# Predicting 3D Chromatin Interactions Using Transformer-Enhanced Deep Learning Models

**DOI:** 10.1101/2025.04.10.647995

**Authors:** Kexin Xu, Li Shen

## Abstract

The three-dimensional (3D) structure of the human genome is essential for regulating gene expression and cellular functions. Chromatin interactions bring distant genomic regions into physical contact, enabling processes like gene regulation, DNA replication, and repair. Disruptions in this organization can lead to diseases such as cancer and genetic disorders. In this study, we propose a Transformer-based deep learning model to predict the chromatin interactions from DNA sequences. By developing a streamlined and efficient data pipeline to handle the sparse and noisy high-throughput chromosome conformation capture (Hi-C) sequencing data, our approach improves both data processing speed and model performance. The Transformer’s ability to capture long-range interactions among genomic regions via attention mechanism, combined with nucleotide position encoding, enables more accurate predictions than purely convolution-based models. This work highlights the potential of Transformer-based network architectures to advance our understanding of genome organization and paves the way for future research with large datasets and advanced network designs.

## 1 Introduction

The three-dimensional (3D) organization of the human genome plays a crucial role in gene regulation, cellular processes, and disease mechanisms. Chromatin interactions, defined as physical contacts between distant genomic regions, facilitate essential processes such as gene expression, DNA replication, and repair [1][2]. Disruptions in chromatin organization are implicated in diseases like cancer [3] and genetic disorders [4], highlighting the need to understand the spatial arrangement of the genome within the cell nucleus. Hi-C experiments [5] provide genome-wide maps of chromatin interactions, offering valuable insights into genome folding. However, these methods are resource-intensive, costly, and limited in scalability, motivating the development of computational approaches to predict chromatin interactions directly from DNA sequences. Early efforts using convolutional neural networks (CNNs), such as Akita [6] and DeepC [7], demonstrated the potential of sequence-based models for predicting chromatin interactions. However, CNNs are inherently limited in their ability to capture long-range dependencies due to the size of the receptive fields, restricting their effectiveness in modeling hierarchical chromatin organization, such as enhancer-promoter loops and topologically associating domains (TADs).

Transformer-based architectures, originally developed for natural language processing [8], have emerged as a powerful solution for modeling long-range dependencies through self-attention mechanisms. In genomics, models like Enformer [9] have demonstrated the potential of Transformers to improve the prediction of gene expression by integrating DNA sequence information across nearly 200Kb regions. Recent advancements, including the DNABERT-2 [10] and the Nucleotide Transformer [11], have further showcased the Transformer’s ability to learn contextual relationships in long DNA sequences. Despite these successes, their applications to chromatin interaction prediction remains underexplored.

To address this gap, we propose DeepChromI, a Transformer-based method designed to predict chromatin interaction matrices using DNA sequences. DeepChromI introduces several key innovations: 1. Training on the largest Hi-C dataset to date: we utilize 117 experiments across 39 cell lines, significantly expanding beyond prior works that used only 1–5 Hi-C experiments. 2. Transformer-based architecture with domain-specific adaptations: our model incorporates novel hierarchical positional encodings to effectively capture long-range chromatin interactions. 3. Scalable and biologically relevant predictions: DeepChromI provides biologically meaningful insights into genome organization, gene regulation, and potentially disease mechanisms while addressing scalability challenges across diverse cell lines. Our work bridges the gap between computational predictions and experimental data, advancing the state of the art in predictive genomics and offering a scalable alternative to Hi-C experiments.

## 2 Methodology

### 2.1 Data Processing

The data processing workflow prepares the input and target for the model: the linear genomic sequences as input and the chromatin interaction matrices as target. Our processing pipeline ensures standardized and high-quality data are produced for model training and evaluation.

#### 2.1.1 Genomic Region Selection and DNA Sequence Processing

The workflow begins by selecting specific regions on the genome as defined in a BED file. In this study, the same BED files from the Enformer paper [9] are used. Each entry in a BED file indicates a genomic interval with a chromosome name, start and end positions. These intervals act as anchors for extracting both DNA sequences and Hi-C interaction data. The DNA sequences are retrieved from the hg38 reference genome. If an interval extends beyond the chromosomal boundaries, the sequence is padded with “N”s to ensure equal sequence length across the dataset. We follow the same train-validation-test split as in the Enformer paper. The length of each interval is modified to be 200Kb using the same center as the original interval to be compatible with the resolution of Hi-C data, which is typically 1Kb, 5Kb or 10Kb. The partitioning yields 34,021 samples for the training set; 2,213 samples for the validation set; and 1,937 samples for the test set.

Each sequence contains four possible nucleotide bases: (A, C, G, T) and N to represent missing value. We use one-hot encoding to transform the sequences into numerical vectors so that A=[1,0,0,0], C=[0,1,0,0], G=[0,0,1,0], T=[0,0,0,1], and N=[0.25,0.25,0.25,0.25]. This transformation yields fixed-size matrices suitable for model input.

#### 2.1.2 Hi-C Interaction Matrix Processing

Hi-C experiments capture the 3D structure of the genome by measuring how frequently different regions of the genome come into physical contact. These interaction frequencies are represented as matrices, where rows and columns correspond to genomic positions, and values indicate the strength of interactions. In this study, a diverse and comprehensive Hi-C dataset comprising 117 high-quality experiments spanning 39 distinct cell lines from the 4D Nucleome (4DN) database [12] is assembled. This represents the largest publicly available Hi-C dataset to date. Only 4DN samples with high interaction counts, excellent coverage, and 1Kb resolution are selected to ensure data quality. The collection includes widely studied cell lines such as GM12878, K562, HepG2, and H1-ESC, as well as primary and differentiated cells. This breadth enables our model to learn both cell-type-invariant chromatin organizations and cell-type-specific interactions. All datasets are processed using the same normalization method to ensure comparability across different experiments.

The interaction matrices corresponding to the genomic regions defined in the BED files are extracted using the hictk package [13]. Smaller matrices are padded with zeros to ensure uniform matrix dimensions. Missing or invalid values are replaced with zeros. Since the interaction matrices are symmetric, about half of the entries are redundant. To reduce memory usage and computational overhead, only the upper triangular portion of each matrix is extracted, including the main diagonal. It is then flattened into a 1D vector. This approach ensures that the model uses only unique chromatin interactions, making training more efficient.

#### 2.1.3 Normalization and Bias Correction

Hi-C data typically contain biases arising from factors such as sequencing depth, restriction enzyme efficiency, and genomic sequence features (e.g., GC content, mappability). These biases are addressed through normalization, primarily using the VC_SQRT (Square Root of Vanilla Coverage) method, as described in Rao et al., 2014 [14]. This approach normalizes matrix values by dividing each entry by the square root of the product of row and column sums, effectively balancing the matrix and reducing sequencing depth biases. By applying VC_SQRT normalization, the influence of high-coverage regions is mitigated, enabling more accurate detection of chromatin interactions and downstream analysis of 3D genome organization.

#### 2.1.4 Handling Distance-Dependent Effects

A notable characteristic of Hi-C data is the high interaction frequency along the matrix diagonal, representing short-range genomic interactions. While biologically meaningful, these strong diagonal signals can overshadow long-range interactions. A diagonal offset approach is implemented to exclude two diagonals near the main diagonal from analysis. This technique helps to focus a model’s attention on more biologically relevant long-range interactions and prevent fitting to trivial short-range signals.

#### 2.1.5 Multi-cell Training Strategy

A key innovation in this study is the multi-cell training framework. Rather than training separate models for each cell line, all cell lines are processed simultaneously. For each genomic region, interaction matrices are extracted from all cell lines and stacked together. Models are trained to predict interactions for all cell lines at once. This multi-task learning strategy encourages feature reuse since all outputs share the same model weights and biases.

### 2.2 Model Architecture

DeepChromI integrates convolution, transformer, and residual connection to predict Hi-C interaction matrices from DNA sequences (Fig. 1). The deep neural network processes genomic sequences through multiple stages to capture both local sequence patterns and long-range chromatin interactions. The network starts with an initial 1D convolutional block that processes the input DNA sequences. This block consists of a 1D convolutional layer with an input channel size of 4 (corresponding to the four nucleotides: A, C, G, and T) and an output channel size of 64. The convolution uses an 11×1 kernel with padding to ensure the input sequence length is preserved. The kernel size is chosen to identify DNA sequence motifs. The output is then passed through batch normalization and ReLU activation, followed by a 1D max-pooling layer with a size of 2 to reduce the sequence length and computational cost.

**Fig. 1.**
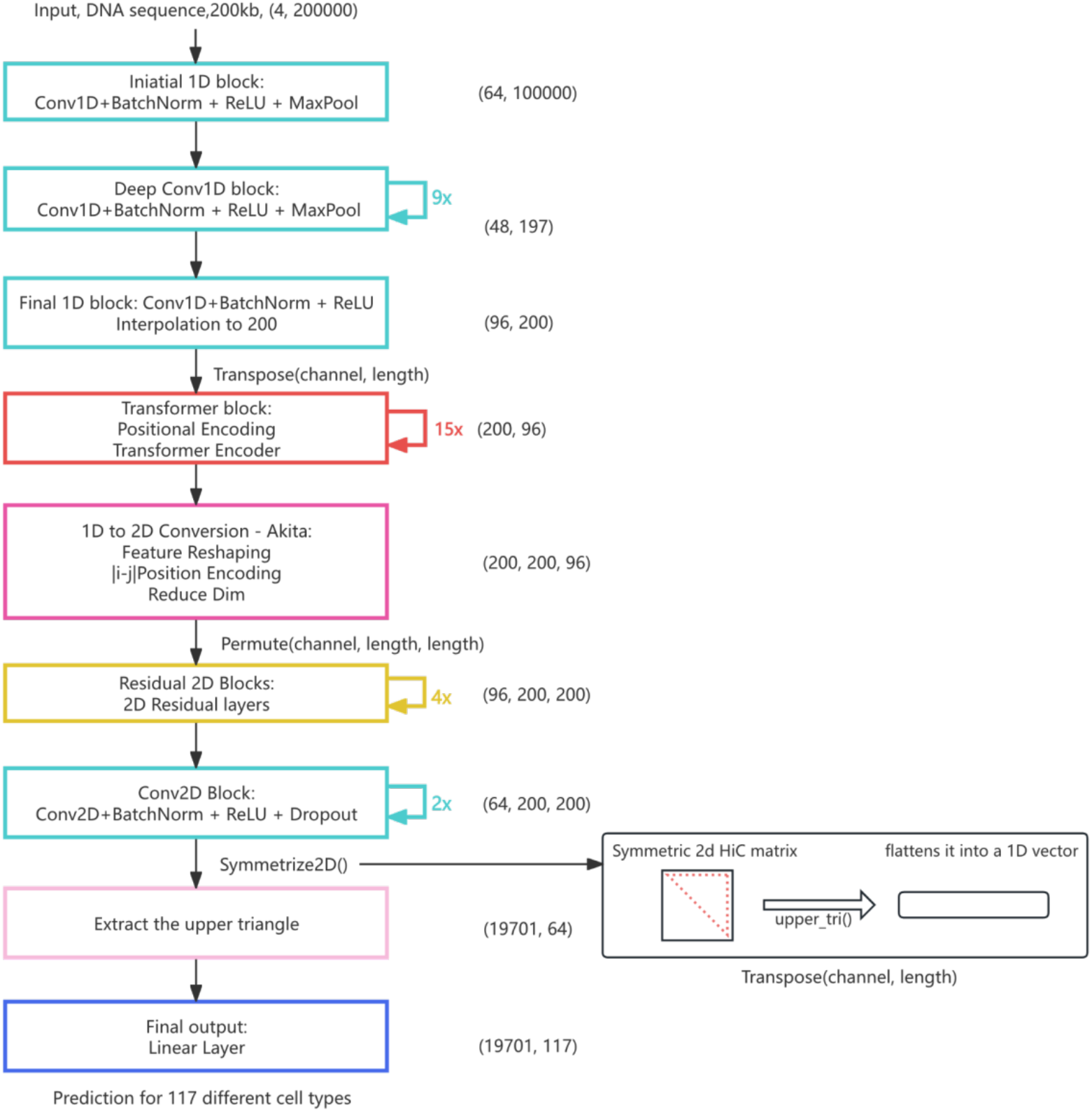
DeepChromI Model Architecture.

After this initial processing stage, the model employs a series of 1D convolutional blocks. Each block consists of a 1D convolutional layer that extracts increasingly complex features. The output channel sizes are defined by a predefined list, alternating between 96 and 48 channels. The convolutional layers use a 3×1 kernel with padding set to 2, allowing the model to learn hierarchical sequence features efficiently. Following each convolutional layer, batch normalization ensures stable training, and a 1D max-pooling layer with a kernel size of 2 progressively reduces the sequence length while retaining the most informative features. a final 1D convolutional block is applied, using a kernel size of 2 with padding set to 1, reducing the feature channels to match the required embedding dimension for the Transformer. This step includes a convolutional layer followed by batch normalization and a ReLU activation function to refine the feature representation. Next, the output undergoes linear interpolation to a fixed length of 200 bins, ensuring consistency in input dimensions for the subsequent Transformer encoder.

The next stage of the model processes the extracted sequence features through a series of transformer layers to refine the representation and capture long-range interactions. To incorporate positional information, positional encoding is added to the interpolated features, which preserves the order of genomic positions within the sequence. The combined features are passed through a multi-layer Transformer encoder, which leverages multi-head self-attention to capture both local and long-range dependencies between distant genomic regions.

Based on the approach introduced by Akita [3], a 1D-to-2D conversion layer is implemented to transform the sequence-based 1D features into a 2D feature map that is suitable for predicting chromatin interactions. This layer computes all pairwise interactions among the genomic bins, converting a (200, 96) tensor into a (200, 200, 96) tensor by averaging the features of all bin pairs. Following the Akita’s design, pairwise positional information (=|i-j|) is incorporated as an additional channel, resulting in a (200, 200, 97) tensor. The subsequent 1×1 convolutional block refines this 2D representation while preserving spatial relationships. It is then processed through multiple 2D convolutional blocks with residual connections, enabling modeling of spatial interactions. Each 2D convolutional block uses a 3×3 kernel size. The first block has 96 channels, while subsequent blocks reduce the output to 64 channels, balancing computational efficiency and representational power. All 2D convolutional blocks use ReLU activation, Batch Normalization, and Dropout for regularization. Because Hi-C interaction matrices are symmetric, a Symmetrize2D layer is added to the last convolutional layer to ensures matrix symmetry by averaging the 2D feature map with its own transposition.

To focus on meaningful long-range interactions, a diagonal offset of 2 is applied to exclude trivial self-interactions near the matrix diagonal. This adjustment directs the model’s attention toward more biologically relevant long-distance chromatin interactions. Furthermore, only the upper triangular portion of the matrix is extracted, reducing redundancy and computational overhead. This design effectively balances precision, biological relevance, and computational efficiency. The final prediction layer consists of linear projection heads that output Hi-C interactions as one 1D vector (size of 19,701) for each of the 117 cell lines, matching our training target format.

Additional architectural designs are implemented to accommodate our data augmentation approach. For data augmentation, two strategies are implemented. First, following Akita’s approach, reverse complement transformation is applied randomly with 50% chance, ensuring strand invariance. Second, random sequence shift up to 11 base pairs is applied to improve positional invariance. A switch reverse triangular layer is used to maintain consistency between transformed sequences and their corresponding Hi-C interactions.

### 2.3 Model Training

The hyperparameters are determined through an automated tuning process using the Optuna [15] package – a popular program for efficient hyperparameter optimization. A search space is defined for key parameters such as the number of Transformer layers and attention heads, dropout rates, and the number of residual and 2D convolutional layers. During this process, Optuna performs multiple trials, where each trial tests a different combination of hyperparameters based on the objective to minimize validation loss. Early stopping is used to prevent overfitting and a learning rate scheduler is adopted for stable convergence. After running 20 trials, the best configuration is automatically selected based on the lowest validation loss achieved. This process results in the final model architecture with 15 Transformer layers, four attention heads, a dropout rate of 0.28, two 2D convolutional layers, and four residual layers (Fig. 1).

For the training process, a fixed learning rate of 0.0001 coupled with a weight decay of 1e-4 are used. To address potential gradient instability issues, gradient clipping with a maximum norm of 1.0 is implemented. The AdamW optimizer is chosen for its ability to handle adaptive learning rates while incorporating weight decay regularization. Mean Squared Error (MSE) is used as the loss function. To enhance the training dynamics, a cosine annealing warm restart scheduler is implemented. This approach allows dynamic learning rate adjustments throughout the training process, helping to navigate the loss landscape more effectively than constant learning rate. The scheduler is proved to be particularly useful in avoiding local minima and promoting better convergence. During the initial three epochs, a warm up phase is incorporated to gradually increase the learning rate, which helps to establish stable training dynamics. A maximum of 16 training epochs is set but the training would halt if the validation loss shows no improvement for 5 consecutive epochs. The batch size is set to be 32.

Several key metrics are monitored on the validation set: these include the MSE, Pearson Correlation Coefficient (R), and R^2^ (Coefficient of Determination). Model checkpoints based on validation loss are saved, ultimately selecting the model that achieves the lowest validation loss for our final evaluation.

## 3 Results

In this section, the performance of the CNN-based Akita [3] model and the Transformer-based DeepChromI model is compared. Both models are trained using the same data pipeline and methodology on 117 Hi-C datasets, spanning 39 distinct cell lines/tissues. The results are analyzed in terms of MSE, R and R^2^ on the test set, highlighting the improvements achieved through the integration of Transformer layers and multi-cell learning.

### 3.1 Model Performance Across Cell Lines/Tissues

Fig. 3 Visualization of the predicted and target Hi-C Interaction Matrices. The heatmaps depict a specific region of Chromosome 11: 75,617,088-75,944,288. The left panel displays the predictions generated by DeepChromI, illustrating its prediction of genomic interactions, while the right panel shows the actual interaction data obtained from the experiment.

The comparison of model metrics between Akita and DeepChromI is summarized in Table 1. DeepChromI consistently outperforms Akita in all metrics. DeepChromI demonstrates a reduced MSE of 1.12, compared to 1.15 for Akita. This indicates that our model can better fit the data and generalize across diverse cell lines/tissues. The average R is improved from 0.78 for Akita to 0.79 for DeepChromI, indicating stronger alignment between predicted and actual interaction matrices across multiple cell lines/tissues. Our model also achieves an average R^2^ value of 0.62, outperforming Akita’s R^2^ of 0.61, suggesting a better overall ability to explain the variance in the data.

**Table 1.** Average results across 117 Hi-C datasets comparing DeepChromI vs. Akita.

| Metric | Akita | DeepChromI |
| --- | --- | --- |
| MSE | 1.15 | 1.12 |
| R (Pearson) | 0.78 | 0.79 |
| $R^2$ | 0.61 | 0.62 |

### 3.2 Cell-Specific Performance

Substantial variation is observed in prediction performance across different cell lines/tissues (Fig. 2). HFFc6 shows the highest prediction accuracy with a R of 0.79, while K562 shows the lowest accuracy with a R of 0.12. Other high-performing cell lines include H9-derived cardiac cells: R of 0.70-0.72 and CyT49 endoderm: R of 0.69.

**Fig. 2.**
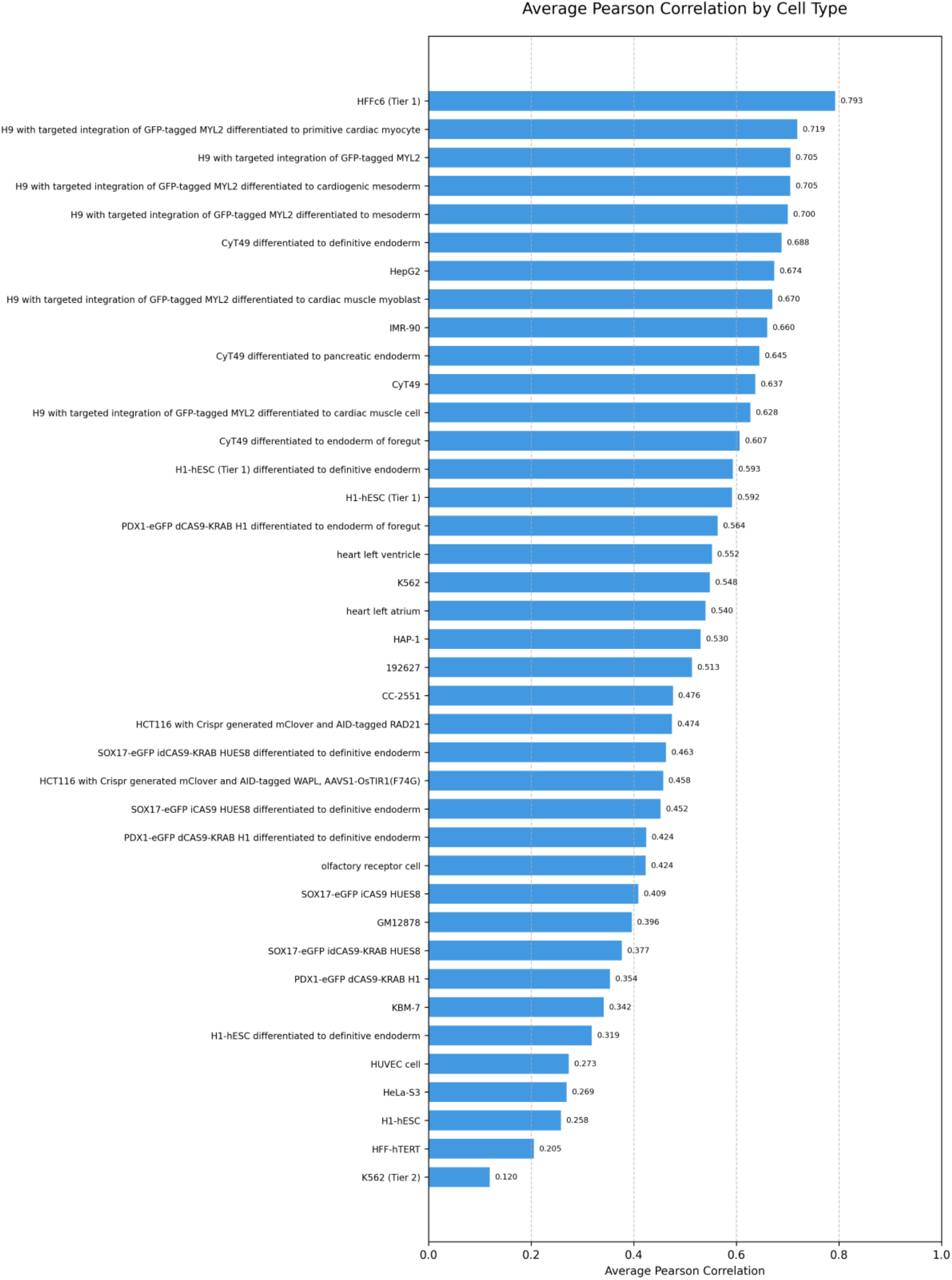
Pearson correlation (R) across different cell lines/tissues.

### 3.3 Visualization of Predicted and Target Hi-C Interaction Matrices

The comparison of predicted and target Hi-C interaction matrices is visualized in Fig. 3, showing an example from the GM12878 cell line (4DNFI1UEG1HD). The Transformer-based DeepChromI successfully captures both local (near-diagonal) and long-range (off-diagonal) interactions, demonstrating its ability to model 3D genome organization effectively.

**Fig. 3.**
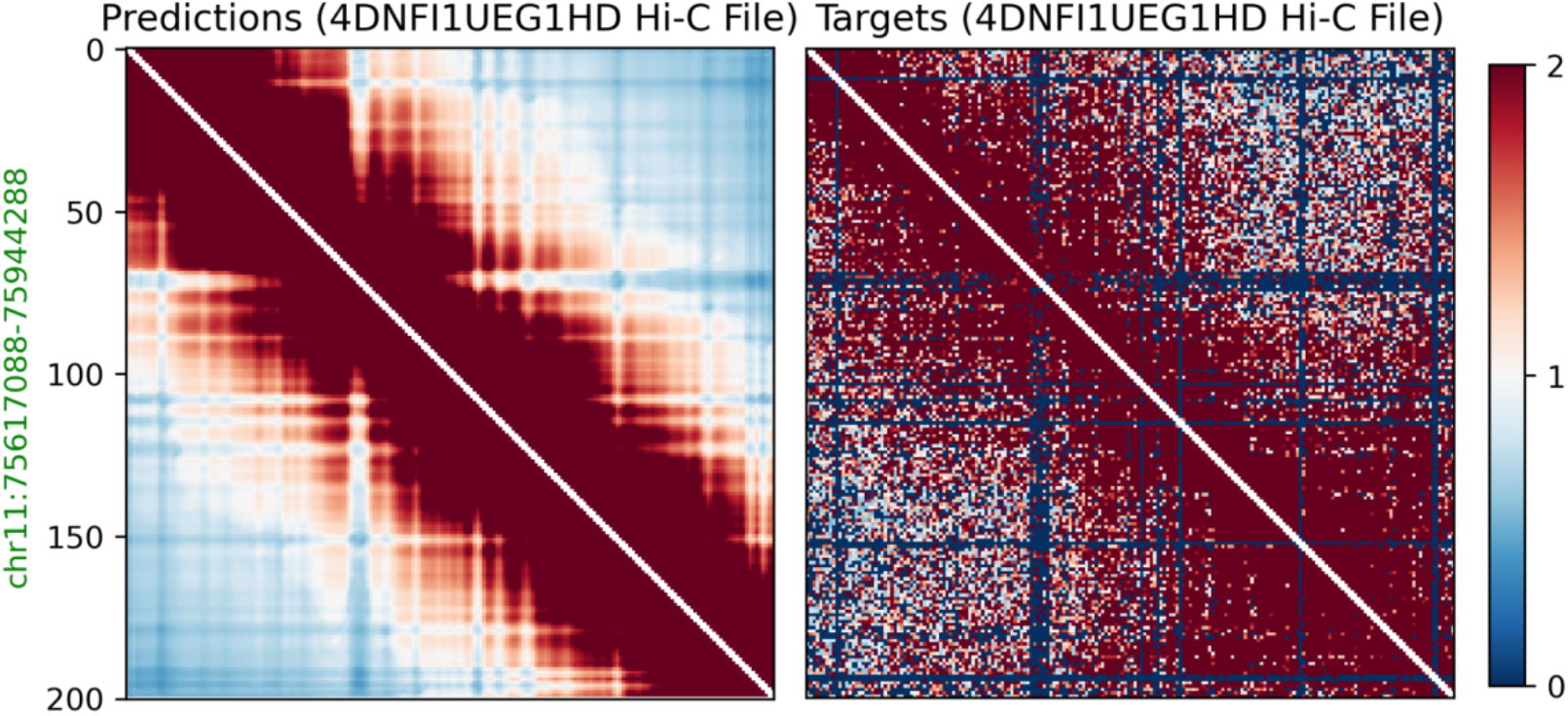
Visualization of the predicted and target Hi-C Interaction Matrices. The heatmaps depict a specific region of Chromosome 11: 75,617,088-75,944,288. The left panel displays the predictions generated by DeepChromI, illustrating its prediction of genomic interactions, while the right panel shows the actual interaction data obtained from the experiment.

## 4 Conclusion

DeepChromI advances the prediction of 3D genome organization by leveraging the Transformer architecture and multi-cell training on a large database of 117 Hi-C datasets. The model achieves superior performance over a CNN-based model. Our result also reveals large performance variation across 39 cell lines/tissues. The Transformer’s self-attention mechanism effectively captures long-range chromatin interactions, while our multi-cell training approach enables the learning of both universal and cell-specific chromatin organizations. These improvements enhance our understanding of the sequence-to-structure relationships in the genome and can help to establish a foundation for investigating the impact of genetic variations on 3D genome organization in diseases.

## Acknowledgement

This work was supported in part through the Minerva computational and data resources and staff expertise provided by Scientific Computing and Data at the Icahn School of Medicine at Mount Sinai and supported by the Clinical and Translational Science Awards (CTSA) grant UL1TR004419 from the National Center for Advancing Translational Sciences.

